# MARCH8 inhibits viral infection by two different mechanisms

**DOI:** 10.1101/2020.04.09.035204

**Authors:** Yanzhao Zhang, Takuya Tada, Seiya Ozono, Satoshi Kishigami, Hideaki Fujita, Kenzo Tokunaga

**Author notes:** Correspondence: Kenzo Tokunaga, Department of Pathology, National Institute of Infectious Diseases, Shinjuku-ku, Tokyo 162-8640, Japan. These authors contributed equally to this work (Yanzhao Zhang, Takuya Tada). Dept. of Microbiology, New York University School of Medicine, New York, New York (Takuya Tada).

## Abstract

Membrane-associated RING-CH 8 (MARCH8) inhibits infection with both HIV-1 and vesicular stomatitis virus G-glycoprotein (VSV-G)-pseudotyped viruses by reducing virion incorporation of envelope glycoproteins. The molecular mechanisms by which MARCH8 targets envelope glycoproteins remain unknown. Here, we show two different mechanisms by which MARCH8 inhibits viral infection. Viruses pseudotyped with the VSV-G mutant, in which cytoplasmic lysine residues were mutated, were insensitive to the inhibitory effect of MARCH8, whereas those with a similar lysine mutant of HIV-1 Env remained sensitive to it. Indeed, the wild-type VSV-G, but not its lysine mutant, was ubiquitinated by MARCH8. Furthermore, the MARCH8 mutant, which had a disrupted cytoplasmic tyrosine motif that is critical for intracellular protein sorting, did not inhibit HIV-1 Env-mediated infection, while it still impaired infection by VSV-G-pseudotyped viruses. Overall, we conclude that MARCH8 reduces viral infectivity by downregulating envelope glycoproteins through two different mechanisms mediated by a ubiquitination-dependent or tyrosine motif-dependent pathway.

## INTRODUCTION

Membrane-associated RING-CH (MARCH) 8 is one of 11 members of the MARCH family of RING-finger E3 ubiquitin ligases, which consist of an N-terminal cytoplasmic tail (CT) domain containing a C4HC3 RING finger (RING-CH finger) motif, two transmembrane (TM) domains, between which a short ectodomain is located, and a C-terminal CT domain^1,2^. MARCH8 downregulates a variety of cellular transmembrane proteins, such as MHC-II^3^, CD86^4^, CD81^5^, CD44^6^, TRAIL receptor 1^7^, CD98^6^, IL-1 receptor accessory protein^8^, and transferrin receptor^9^. We have recently reported that MARCH8 reduces HIV-1 infectivity by downregulating HIV-1 envelope glycoproteins (Env) from the cell surface, resulting in a reduced incorporation of Env into virions^10^. Intriguingly, vesicular stomatitis virus G-glycoprotein (VSV-G) was even more sensitive to the inhibitory effect of MARCH8. In the case of HIV-1 Env, it is retained intracellularly without degradation after cell-surface downregulation. In contrast, VSV-G is not only downregulated from the cell surface but also undergoes lysosomal degradation by MARCH8^10^. In this regard, we hypothesized that VSV-G, whose cytoplasmic tail is lysine-rich (5 out of 29 amino acids), could be readily ubiquitinated by the E3 ubiquitin ligase MARCH8 and therefore undergo lysosomal degradation, whereas HIV-1 Env carries only two lysines (out of 151 amino acids) in its cytoplasmic tail and may rarely undergo degradation after being trapped by MARCH8. In this study, we created lysine mutants of both HIV-1 Env and VSV-G, together with newly generated MARCH8 mutants to explore the hypothesis described above. The results with these mutants show that MARCH8 targets HIV-1 Env and VSV-G by two different inhibitory mechanisms.

## RESULTS AND DISCUSSION

We have recently reported that MARCH8 inhibits lentiviral infection by reducing virion incorporation of both HIV-1 Env and VSV-G in a RING-CH domain-dependent manner. Because the RING-CH domain is known to be essential for the E3 ubiquitin ligase activity of MARCH8, we asked whether these envelope glycoproteins are susceptible to MARCH8-mediated ubiquitination. To investigate this, we first created the VSV-G mutant CT5K/R in which five arginine residues were introduced in place of cytoplasmic lysine residues that could be ubiquitination targets (Fig. 1A, upper). We also generated the HIV-1 Env gp41 mutant CT2K/R harboring two arginines in place of the cytoplasmic lysines (Fig. 1A, lower). Then, we prepared HIV-1 luciferase reporter viruses pseudotyped with the mutant envelope glycoproteins (VSV-G CT5K/R and HIV-1 Env CT2K/R) from 293T cells transiently expressing MARCH8, and compared their viral infectivity with that of control viruses pseudotyped with wild-type (WT) envelope glycoproteins. The infectivity of viruses harboring either VSV-G CT5K/R or HIV-1 gp41 Env CT2K/R was almost comparable to WT-enveloped viruses (Figs. 1B and 1C). As expected, the virus pseudotyped with VSV-G CT5K/R was completely resistant to MARCH8 (Fig. 1B). In contrast, the HIV-1 Env CT2K/R-pseudotyped virus was still susceptible to the inhibitory effect of MARCH8 (Fig. 1C). Consistent with these results, immunofluorescence staining showed that the five lysine mutations in the CT domain of VSV-G conferred resistance to MARCH8-mediated intracellular degradation (Fig. 1D), whereas the two lysine mutations in the CT domain of HIV-1 Env had no effect on its cell-surface downregulation by MARCH8 (Fig. 1E). It should be noted that MARCH8-resistant VSV-G CT5K/R colocalized with MARCH8 (Fig. 1D). We thus speculated that MARCH8 ubiquitinates lysine residues of VSV-G but not of HIV-1 Env at their CT domains.

**Figure 1.**
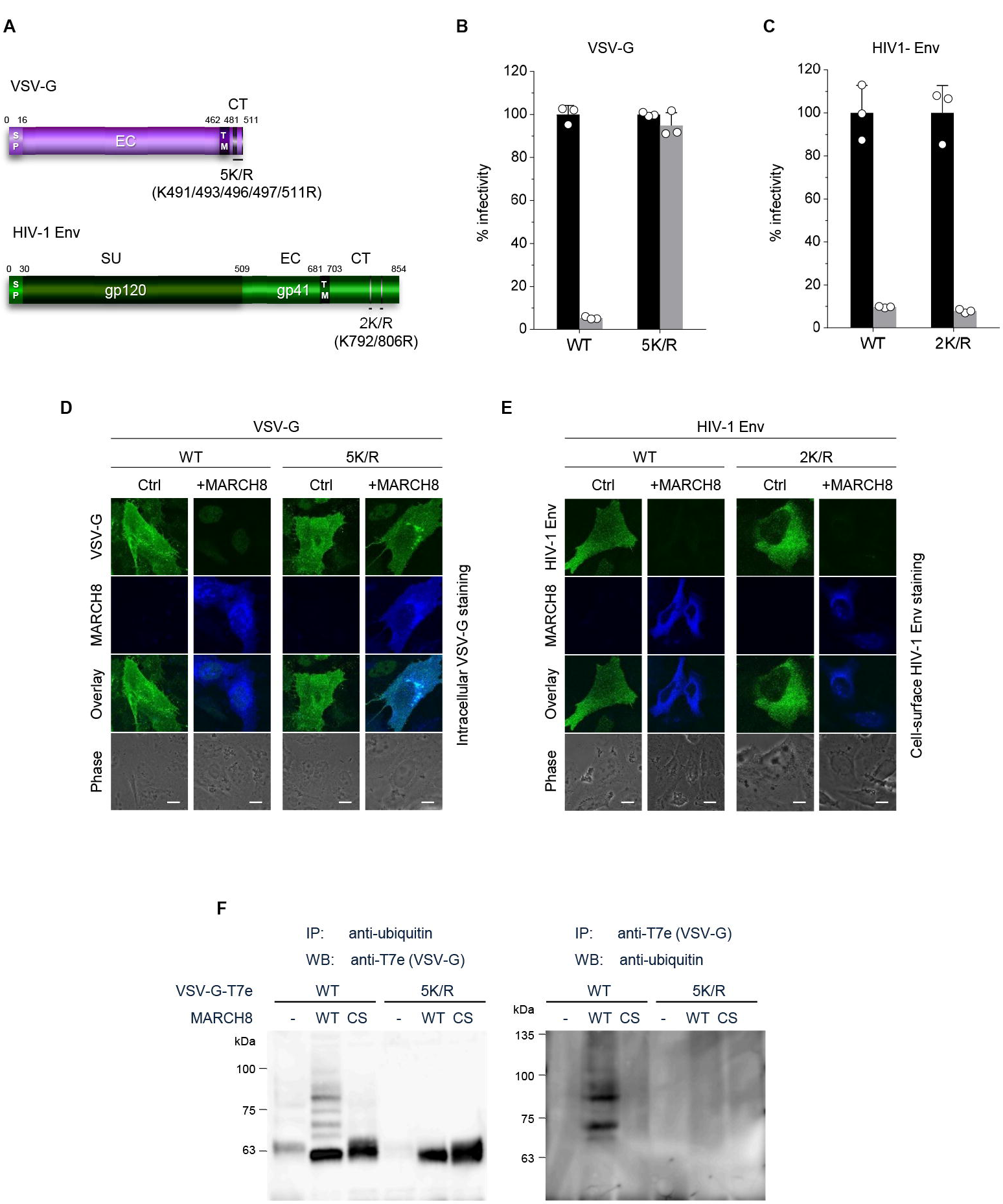
(**A**) Schematic structure of the lysine mutants of VSV-G (CT5K/R; upper) and HIV-1 Env (CT2K/R; lower). SP, signal peptide; EC, extracellular domain; TM, transmembrane domain; CT, cytoplasmic tail; SU, surface subunit. (**B**) Infectivity of viruses prepared from 293T cells cotransfected with Env-defective HIV-1 luciferase (luc) reporter proviral DNA and either the VSV-G wild-type (WT) or CT5K/R mutant plasmid together with either a control (Ctrl) (black) or HA-MARCH8 (gray) plasmid. Data are shown as a percentage of the viral infectivity in the absence of MARCH8 when WT VSV-G was used (mean + s.d. from three independent experiments). (**C**) Infectivity of viruses prepared as shown in *B*, except for using either the WT HIV-1 Env or its CT2K/R mutant plasmid (mean + s.d. from three independent experiments). (**D**) The VSV-G lysine mutant is resistant to MARCH8-mediated intracellular degradation. Shown are immunofluorescence-based analyses of the intracellular levels of either the WT or lysine mutant VSV-G with or without MARCH8 in transfected HOS cells. Scale bars, 10 μm. (**E**) The lysine mutant of HIV-1 Env is still sensitive to MARCH8-induced downregulation from the cell surface. Immunofluorescence images show cell-surface levels of either the WT or lysine mutant HIV-1 Env with or without MARCH8 in transfected HOS cells. Scale bars, 10 μm. (**F**) Lysine residues at the CT domain of VSV-G are ubiquitinated by MARCH8. The ubiquitination of the WT or lysine mutant VSV-G tagged with T7-epitope (T7e) in cells expressing control or MARCH8 (WT or RING-CH mutant (CS)) was examined by immunoprecipitation of either ubiquitinated proteins with an anti-ubiquitin antibody (left panel) or of T7e-tagged VSV-G with an anti-T7e antibody (right panel), followed by immunoblotting with an antibody to either T7e (left panel) or ubiquitin (right panel), respectively.

To investigate this possibility, we performed immunoprecipitation (IP)/Western-based ubiquitination assays. In cells coexpressing WT HA-MARCH8, VSV-G was efficiently ubiquitinated, whereas in cells coexpressing the RING-CH mutant of HA-MARCH8, the ubiquitination of VSV-G was lost. More importantly, the VSV-G lysine mutant CT5K/R did not undergo MARCH8-mediated ubiquitination, as expected (Fig. 1F), suggesting that the lysine residues at the CT domain of VSV-G are specifically ubiquitinated by MARCH8. These findings are consistent with the immunofluorescence results (Figs. 1D and 1E). We therefore conclude that lysine residues at the CT domain of VSV-G are ubiquitinated by MARCH8, which determines the difference in the MARCH8-mediated intracellular fate between these viral glycoproteins.

Because unlike VSV-G, HIV-1 Env was retained intracellularly without degradation, as we previously reported^10^, we hypothesized that these viral envelope glycoproteins might undergo endocytosis with different mechanisms of action. It has been reported that another MARCH family member, MARCH11, has a conserved tyrosine-based motif, YXXϕ, which is known to be recognized by the adaptor protein (AP) μ-subunits in the C-terminal CT domain^11^. We therefore looked for the same motif(s) in MARCH8 and found the ^222^YxxL^225^ and ^232^YxxV^235^ sequences in the CT domain at the C-terminus (Fig. 2A). Based on this finding, we generated tyrosine motif mutants of MARCH8, in which either ^222^Y or ^232^Y was mutated to alanine (designated ^222^AxxL^225^ or ^232^AxxV^235^). The protein expression in cells transfected with each MARCH8 plasmid was confirmed by immunoblotting using an anti-hemagglutinin (HA) antibody (Fig. 2B). Then, we examined whether these YXXϕ motifs are important for the antiviral activity of MARCH8. The infectivity of VSV-G–pseudotyped viruses was still inhibited by the expression of both ^222^AxxL^225^ and ^232^AxxV^235^ MARCH8 mutants (Fig. 2C). In contrast, the infectivity of HIV-1 Env– pseudotyped viruses was not impaired by ^222^AxxL^225^ MARCH8 expression but was reduced by that of ^232^AxxV^235^ and WT MARCH8 (Fig. 2D), suggesting that the first tyrosine motif (^222^YxxL^225^) is involved in the antiviral activity of MARCH8 against HIV-1 Env but not VSV-G. We previously reported that the MARCH8-mediated reduction in viral infectivity was due to reduced entry efficiency, resulting from a decreased virion incorporation of envelope glycoproteins through their downregulation from the cell surface^10^. Therefore, by performing a β-lactamase (BlaM)-fused viral protein R (Vpr)-based entry assay, we first focused on whether the loss of function of ^222^AxxL^225^ MARCH8 against HIV-1 Env but not VSV-G would indeed be due to the loss of its inhibitory activity on viral entry. Whereas the entry of VSV-G–pseudotyped HIV-1 prepared from cells expressing either WT or ^222^AxxL^225^ MARCH8 was reduced compared with that of the control virus (Fig. 2E), the inhibition of the entry of whole HIV-1 virions was abrogated in viruses produced from cells expressing ^222^AxxL^225^, as expected (Fig. 2D). We further analyzed whether this motif of MARCH8 is indeed involved in the reduced virion incorporation of HIV-1 Env, which results from its cell-surface downregulation. To address this, we conducted flow cytometric analysis and quantified the levels of cell-surface and intracellular expression of Env glycoproteins. In accordance with the results obtained in infectivity assays (Fig. 2C), ^222^AxxL^225^ MARCH8 still reduced intracellular VSV-G expression as well as WT MARCH8 did (Fig. 2G). On the other hand, the mutant MARCH8 had a completely abrogated ability to downregulate cell-surface HIV-1 Env, whereas WT expression led to the downregulation of HIV-1 Env from the cell surface and its intracellular retention (Fig. 2H), as we previously observed^12^. The results were consistent with those of the inhibitory activity of MARCH8 against the virion incorporation of HIV-1 Env (Fig. 2I). We therefore conclude that ^222^YxxL^225^ is critical for the MARCH8-mediated downregulation of HIV-1 Env but not VSV-G, which results in its reduced virion incorporation leading to impaired viral entry.

**Figure 2.**
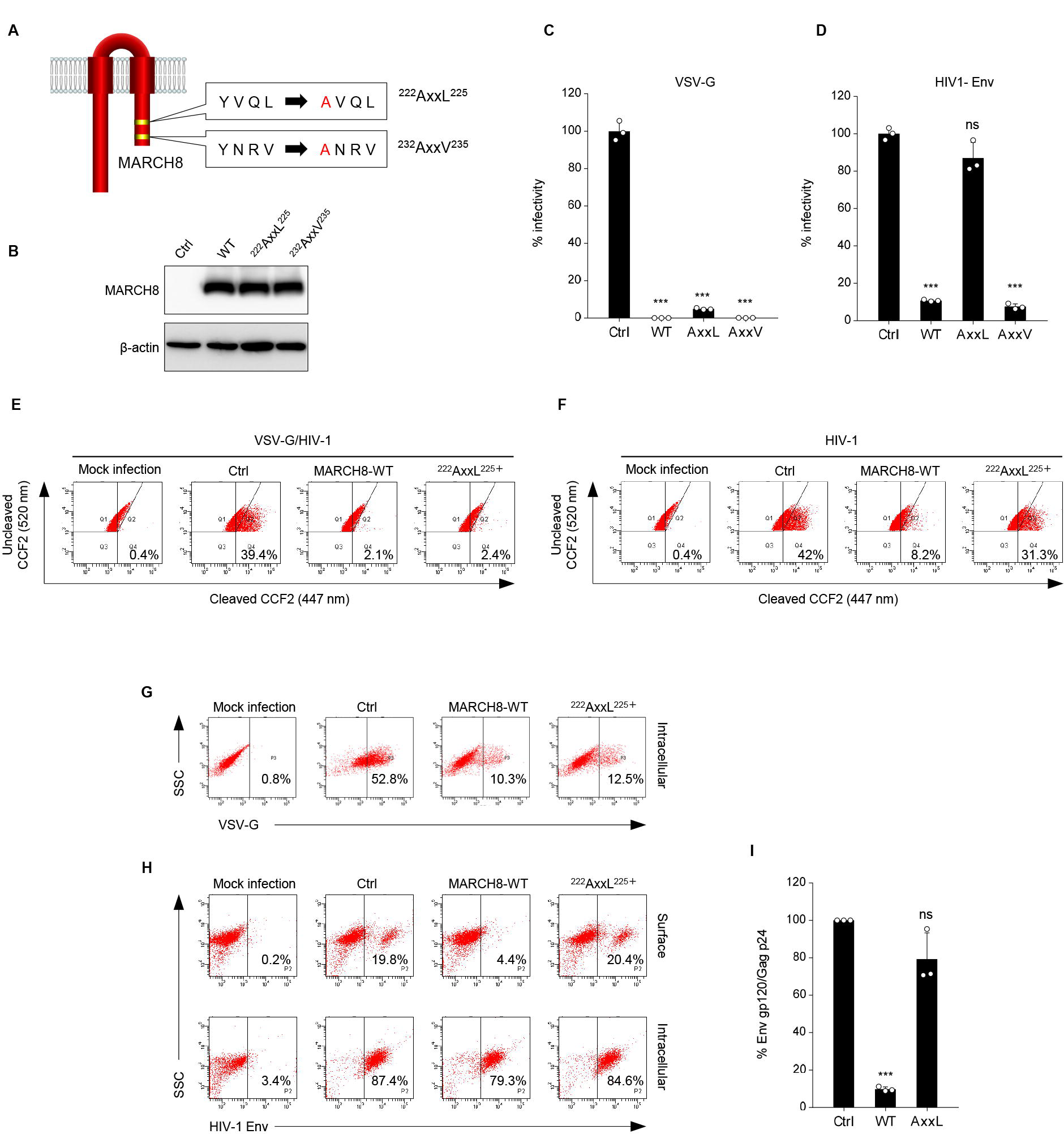
(**A**) Schematic structure of YxxF motif mutants of MARCH8 (^222^YxxL^225^ and ^232^YxxV^235^). (**B**) Western blot analysis was performed by using extracts from 293T cells transfected with HA-tagged MARCH8 expression plasmids. Antibodies specific for HA were used to detect MARCH8 proteins. (**C, D**) Infectivity of viruses prepared from 293T cells cotransfected with Env-defective HIV-1 luciferase (luc) reporter proviral DNA and either a control (Ctrl), HA-WT, HA-^222^AxxL^225^ or HA-^232^AxxV^235^ MARCH8 plasmid, together with either (**C**) the VSV-G expression plasmid or (**D**) the HIV-1 Env expression plasmid. Data are shown as a percentage of the viral infectivity in the absence of MARCH8 (mean + s.d. from three independent experiments). ns; \*\*\**P* < 0.0005 compared with the Ctrl using two-tailed unpaired *t*-tests. (**E, F**) BlaM-Vpr-based viral entry assay using VSV-G-pseudotyped viruses (**E**) or NL4-3 whole viruses (**F**) produced from cells expressing either control, WT MARCH8, or the ^222^AxxL^225^ mutant. Representative FACS dot plots are shown from four independent experiments. (**G**) VSV-G is downregulated by both WT and ^222^AxxL^225^ mutant MARCH8, (**H**) whereas the cell-surface expression of HIV-1 Env is not affected by the mutant MARCH8. (**I**) ^222^AxxL^225^ MARCH8 expression in producer cells is unable to decrease HIV-1 gp120 levels in viral supernatants. ELISA-based levels of Env gp120 in viral supernatants from 293T cells cotransfected with luc reporter proviral DNA and NL-Env plasmid, together with either MARCH8 WT or its ^222^AxxL^225^ mutant. Representative data from three independent experiments are shown as percent gp120 Env/p24 Gag in the supernatants relative to that from control cells. (mean + s.d. from three independent experiments). ns; \*\*\**P* < 0.0005 compared with the Ctrl using two-tailed unpaired *t*-tests.

In summary, we first show the two different mechanisms by which MARCH8 inhibits viral infections, one being a ubiquitin-dependent downregulation that mediates lysosomal degradation of VSV-G whose cytoplasmic lysine residues are recognized by the RING-CH domain of MARCH8 (Fig. 3, left), and the other, a YxxΦ motif-dependent downregulation that could explain the intracellular retention of HIV-1 Env without degradation after cell-surface downregulation (Fig. 3, right). In terms of the latter mechanism, although this could be attributed to the AP-dependent trafficking that requires the YxxΦ motif, which binds to AP μ-subunits, our preliminary results showed no interaction between these subunits and MARCH8. We have recently reported that MARCH1 and MARCH2 are also antiviral MARCH family members that inhibit HIV-1 infection, although their antiviral activity is less robust, which is probably due to the lower protein expression and/or stability than that of MARCH8^12^. Because MARCH1 and MARCH2 also harbor the Yxxϕ motif in their C-terminal CT domains, it would be intriguing to verify the importance of the motif in these proteins. Overall, our present findings are consistent with our previous studies showing that MARCH8-induced downregulation of VSV-G leads to lysosomal degradation, while that of HIV-1 Env results in intracellular retention without degradation. Thus, we conclude that MARCH8 targets HIV-1 Env and VSV-G by two different inhibitory mechanisms (either ubiquitin-dependent or YxxF motif-dependent downregulation). Further investigations will clarify the more detailed host defense mechanisms of this protein.

**Figure 3.**
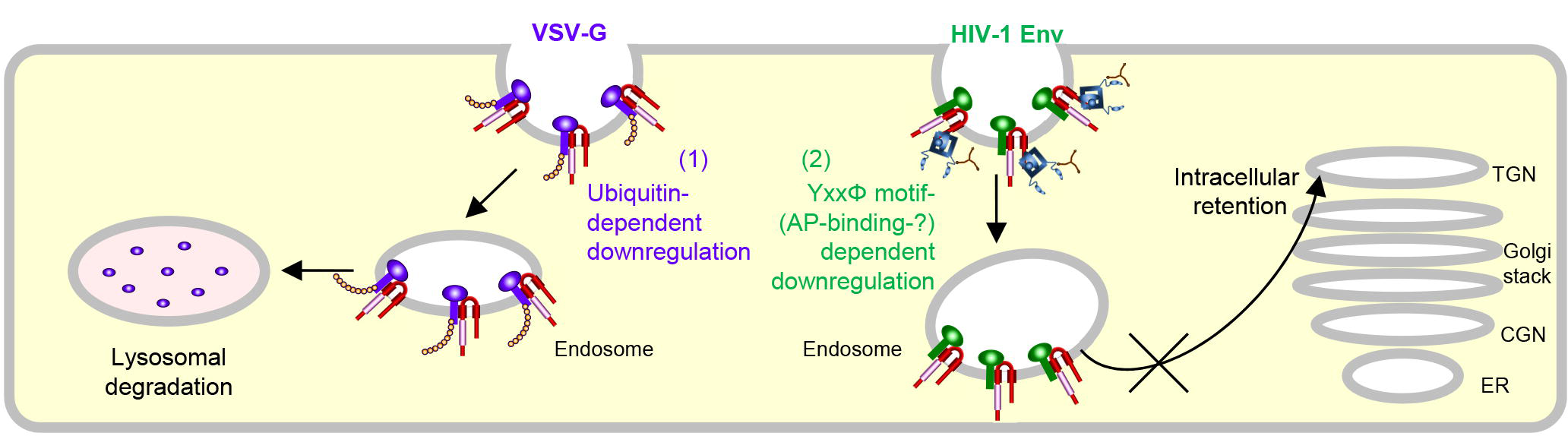
Schematic diagram of two different molecular mechanisms by which MARCH8 inhibits viral infection. *Left*, MARCH8 (red) downregulates VSV-G (violet) in a ubiquitin-dependent manner. The RING-CH domain (pink) of MARCH8 recognizes VSV-G’s cytoplasmic lysine residues, which results in ubiquitin conjugation (shown as orange beads), leading to lysosomal degradation; *Right*, MARCH8 downregulates HIV-1 Env (green) in a YxxF motif-dependent manner. The tyrosine motif located in the C-terminal CT of MARCH8 likely interacts with the adaptor protein μ-subunits (navy) (if this is the case with μ2 or μ1, clathrin (brown) is involved in this step), resulting in the intracellular retention of HIV-1 Env without degradation. It should be noted that the downregulation of these viral glycoproteins might not necessarily occur at the plasma membrane. The nucleus and other organelles are not shown.

## METHODS

### DNA constructs

The Env-deficient HIV-1 proviral indicator construct pNL-Luc2-E(-), HIV-1 Gag-Pol expression plasmid pC-GagPol-RRE, HIV-1 Env-expression vector pC-NLenv, HIV-1 Rev expression plasmid pCa-Rev, VSV-G expression plasmid pC-VSVg and its C-terminally T7-epitope-tagged version pVSVg-T7E, GFP expression plasmid pCa-EGFP, Vpr/β-lactamase (BlaM) expression plasmid pMM310, and MARCH8 expression plasmid either pC-MARCH8 or pC-HA-MARCH8 and its RING-CH mutant pC-HA-MARCH8-CS have previously been described elsewhere^10^. The HIV-1 Env gp41 mutant CT2K/R, in which cytoplasmic lysine residues at positions 792 and 806 were mutated to arginine, and the VSV-G mutant CT5K/R, in which cytoplasmic lysine residues at positions 491, 493, 496, 497, and 511 were mutated to arginine residues, were created by inserting overlapping PCR fragments into *Mfe*I/*Xho*I-digested pC-NLenv and by inserting PCR fragments into *Kpn*I/*Not*I-digested pC-VSVg (or pVSVg-T7E), respectively. The MARCH8 mutant ^222^AxxL^225^ or ^232^AxxV^235^, in which a tyrosine residue at position 222 or 232 was mutated to an alanine residue, was generated by inserting overlapping PCR fragments into the *Kpn*I/*Xho*I-digested pCAGGS or *Xho*I-*Not*I-digested pCAGGS-NHA to create an untagged or N-terminally HA-tagged expression plasmid. All constructs were verified by a DNA sequencing service (FASMAC).

### Cell maintenance and generation of stable cell lines

293T, MT4, HeLa, MAGIC5 (HeLa derivative^13^), and HOS cells were maintained under standard conditions. Cells were originally obtained from ATCC (except MAGIC5 cells) and routinely tested negative for mycoplasma contamination (PCR Mycoplasma Detection kit, Takara).

### Virion production and infectivity assays

To prepare VSV-G-pseudotyped or HIV-1-Env-pseudotyped luciferase reporter viruses, 2.5 × 10^5^ 293T cells were cotransfected with 120 ng of the MARCH expression plasmid (WT or RING-CH mutants), 20 ng of pC-VSVg, pC-VSVg-CT5K/R, pC-NLenv, or pC-NLenv-CT2K/R, 500 ng of pNL-Luc2-E(-), and an empty vector up to 1 μg of total DNA, using FuGENE 6 (Promega). Sixteen hours later, the cells were washed with phosphate-buffered saline, and 1 ml of fresh complete medium was added. After 24 h, supernatants were treated as described above and then harvested. The p24 antigen levels in viral supernatants were measured by an HIV-1 p24 antigen capture ELISA (XpressBio). Transfection efficiencies were normalized to the luciferase activity. To determine viral infectivity, 1 × 10^4^ MAGIC5 cells were incubated with 1 ng of p24 antigen from the HIV-1 supernatants. After 48 h, cells were lysed in 100 μl of One-Glo Luciferase Assay Reagent (Promega). The firefly luciferase activity was determined with a Centro LB960 (Berthold) luminometer.

### Immunoblotting assays

Protein expression of constructs was confirmed by Western blot analyses as described elsewhere^10,12^. Briefly, cells transfected as described above were lysed in 500 μl of lysis buffer containing 1.25% n-octyl-β-D-glucoside (Dojindo), and Complete protease inhibitor cocktail (Roche Applied Science). Cell extracts were then subjected to gel electrophoresis and transferred to a nitrocellulose membrane, followed by probing with an anti-HA mouse monoclonal antibody (Sigma-Aldrich, H9658) or an anti-β-actin mouse monoclonal antibody (Sigma-Aldrich, A5316). Proteins were then visualized by chemiluminescence using an ECL Western blotting detection system (GE Healthcare) and monitored by using a LAS-3000 imaging system (FujiFilm).

### Viral entry assays

A β-lactamase (BlaM)-fused viral protein R (Vpr)-based entry assay was performed as described elsewhere ^10,12^. Briefly, HIV-1 particles containing a fusion protein of Vpr and BlaM-Vpr were produced by cotransfection of 293T cells with pNL4-3, pMM310^14^ encoding BlaM-Vpr, and either pC-MARCH8, pC-MARCH8-^222^AxxL^225^, pC-MARCH8-^232^AxxV^235^, or the control vector. Similarly, VSV-G-pseudotyped HIV-1 particles containing BlaM-Vpr were prepared by cotransfection with pNL-Luc-E(-), pC-VSVg, pMM310, and either pC-MARCH8, pC-MARCH8-^222^AxxL^225^, pC-MARCH8-^232^AxxV^235^, or the control vector. The produced viruses were normalized to the p24 antigen level (100 ng) and used for infection of the CD4^+^ T cell line MT4 (5 × 10^5^ cells) at 37 °C for 4 h to allow viral entry. After extensive washing with Hank’s balanced salt solution (HBSS; Invitrogen), cells were incubated with 1 μM CCF2-AM dye (Invitrogen), a fluorescent substrate of BlaM, in HBSS containing 1 mg ml^−1^ Pluronic F-127 surfactant (Invitrogen) and 0.001% acetic acid for 1 h at room temperature and then washed with HBSS. Cells were further incubated for 14 h at room temperature in HBSS supplemented with 10% FBS, washed three times with PBS and fixed in a 1.2% paraformaldehyde solution. Fluorescence was monitored at 520 and 447 nm by flow cytometry using BD FACS Canto II (BD Bioscience), and the data were collected and analyzed with BD FACS Diva Software (BD Bioscience).

### Env incorporation assays

HIV-1 gp120 ELISA-based Env incorporation assays were performed by using an HIV-1 gp120 antigen capture ELISA kit (Advanced BioScience Laboratories) as described elsewhere^12^.

### Ubiquitination assays

293T cells (5 × 10^5^) were cotransfected with 0.8 μg of pC-VSVg-T7E, 0.2 μg of the empty vector, and 0.8 μg of pC-HA-MARCH8. After 48 h, cells were lysed in TBS-T buffer (50 mM Tris-HCl buffer (pH 7.5), 0.15 M NaCl, 1% Triton X-100, and 0.5% deoxycholic acid) containing a protease inhibitor cocktail and 10 mM N-ethylmaleimide, as an inhibitor of deubiquitination enzymes. The mixture was centrifuged at 21,500 × g for 15 min, and the supernatant was used as total cell lysate for immunoblotting or immunoprecipitation. Fifty microliters of Protein A-coupled Sepharose 4B (GE Healthcare, 17-0780-01) was preincubated for 2 h at 4 °C with 4 μg of the appropriate antibody (anti-T7 epitope rabbit polyclonal antibody, MBL, PM022; anti-ubiquitin mouse monoclonal antibody Clone FK2, Cayman, 14220). Total cell lysate was incubated with antibody-coupled Sepharose for 20 h at 4 °C. The Sepharose was washed three times with TBS-T buffer and one time with PBS before the immunoprecipitated proteins were eluted with SDS sample buffer. To evaluate the ubiquitination states of the immunoprecipitated proteins, proteins immunoprecipitated with an anti-T7 epitope rabbit antibody were subjected to Western blotting with an anti-ubiquitin mouse antibody, whereas proteins immunoprecipitated with the anti-ubiquitin mouse antibody were subjected to Western blotting with the anti-T7 epitope rabbit antibody. Immunoreactive bands were detected using an ECL detection kit (EzWestLumi plus, ATTO) with a ChemiDoc imaging system (Bio-Rad).

### Immunofluorescence microscopy

HOS cells were plated on 13-mm coverslips, cotransfected with the indicated plasmids, 0.5 μg of either pC-NLenv or pC-VSVg-T7E, 0.1 μg of pC-Gag-Pol, 0.05 μg of pCa-Rev, and 0.3 μg of the pC-HA-MARCH8 expression plasmids (WT or CS-mutant) using FuGENE6 and cultured for 24 h. For the total staining of both VSV-G and MARCH8, cells were fixed with 4% paraformaldehyde for 30 min on ice and permeabilized with 0.05% saponin. The fixed cells were incubated with both primary antibodies anti-T7 epitope mouse monoclonal antibody (Novagen, 69522-4) and anti-HA goat polyclonal antibody (GenScript, A00168-40). The secondary antibodies, Alexa 488 donkey anti-mouse IgG (Molecular Probes, A-21202) and Alexa 647 donkey anti-goat IgG (Molecular Probes, A-21447) were used for the double staining assay. For the cell surface staining of gp120 protein, cells were incubated with an anti-gp120 goat polyclonal antibody (Abcam, Ab21179) at 4 °C for 5 min and washed with PBS at 4 °C before fixation. Fixation was performed with 4% paraformaldehyde for 30 min on ice, and fixed cells were permeabilized with 0.05% saponin (Sigma-Aldrich) to detect the intracellular expression and localization of MARCH8 proteins. The coverslips were incubated with the anti-HA mouse monoclonal antibody (Sigma Aldrich) for 1 h, washed with PBS and incubated for 30 min with the secondary antibodies Alexa 488 donkey anti-goat IgG (Molecular Probes, A-11055) and Alexa 647 donkey anti-mouse IgG (Molecular Probes, A-31571) for the double staining assay. Confocal images were obtained with a FluoView FV10i automated confocal laser-scanning microscope (Olympus; Tokyo).

### Statistical analyses

Column graphs that combine bars and individual data points were created with GraphPad Prism version 8.04. *P*-values generated from two-tailed paired *t*-tests for data represented in Figures 2C, 2D, and 2I.

## ACKNOWLEDGMENTS

This work was supported by grants from the Japan Society for the Promotion of Science (KAKENHI, 18K07156) to K.T.

## AUTHOR CONTRIBUTIONS

Y.Z., T.T., S.O., H.F. and K.T. performed the experiments and analyzed the data. Y.Z., H.F. and

K.T. discussed the data. S.K. provided reagents. K.T. conceived the study, supervised the work and wrote the paper. All authors read and approved the final manuscript.

## REFERENCES

1. Bartee, E., Mansouri, M., Hovey Nerenberg, B. T., Gouveia, K. & Fruh, K. Downregulation of major histocompatibility complex class I by human ubiquitin ligases related to viral immune evasion proteins. J Virol. 78, 1109–1120 (2004).

2. Goto, E. et al. c-MIR, a human E3 ubiquitin ligase, is a functional homolog of herpesvirus proteins MIR1 and MIR2 and has similar activity. J Biol Chem. 278, 14657–14668 (2003).

3. Ohmura-Hoshino, M. et al. Inhibition of MHC class II expression and immune responses by c-MIR. J Immunol. 177, 341–354 (2006).

4. Tze, L. E. et al. CD83 increases MHC II and CD86 on dendritic cells by opposing IL-10-driven MARCH1-mediated ubiquitination and degradation. J Exp Med. 208, 149–165 (2011).

5. Bartee, E. et al. Membrane-Associated RING-CH proteins associate with Bap31 and target CD81 and CD44 to lysosomes. PLoS One. 5, e15132 (2010).

6. Eyster, C. A. et al. MARCH ubiquitin ligases alter the itinerary of clathrin-independent cargo from recycling to degradation. Mol Biol Cell. 22, 3218–3230 (2011).

7. van de Kooij, B. et al. Ubiquitination by the membrane-associated RING-CH-8 (MARCH-8) ligase controls steady-state cell surface expression of tumor necrosis factor-related apoptosis inducing ligand (TRAIL) receptor 1. J Biol Chem. 288, 6617–6628 (2013).

8. Chen, R., Li, M., Zhang, Y., Zhou, Q. & Shu, H. B. The E3 ubiquitin ligase MARCH8 negatively regulates IL-1beta-induced NF-kappaB activation by targeting the IL1RAP coreceptor for ubiquitination and degradation. Proc Natl Acad Sci U S A. 109, 14128–14133 (2012).

9. Fujita, H., Iwabu, Y., Tokunaga, K. & Tanaka, Y. Membrane-associated RING-CH (MARCH) 8 mediates the ubiquitination and lysosomal degradation of the transferrin receptor. J Cell Sci. 126, 2798–2809 (2013).

10. Tada, T. et al. MARCH8 inhibits HIV-1 infection by reducing virion incorporation of envelope glycoproteins. Nat Med. 21, 1502–1507 (2015).

11. Morokuma, Y. et al. MARCH-XI, a novel transmembrane ubiquitin ligase implicated in ubiquitin-dependent protein sorting in developing spermatids. J Biol Chem. 282, 24806–24815 (2007).

12. Zhang, Y. et al. Membrane-associated RING-CH (MARCH) 1 and 2 are MARCH family members that inhibit HIV-1 infection. J Biol Chem. 294, 3397–3405 (2019).

13. Mochizuki, N. et al. An infectious DNA clone of HIV type 1 subtype C. AIDS Res Hum Retroviruses. 15, 1321–1324 (1999).

14. Tobiume, M., Lineberger, J. E., Lundquist, C. A., Miller, M. D. & Aiken, C. Nef does not affect the efficiency of human immunodeficiency virus type 1 fusion with target cells. J Virol. 77, 10645–10650 (2003).

